# Benchmarking attention-based methods for vision transformers’ interpretability in retinal fundus imaging

**DOI:** 10.64898/2026.06.15.732470

**Authors:** Sacha Bors, Michael Beyeler, Olga Trofimova, VascX Consortium, David Presby, Dennis Bontempi, Sven Bergmann

**Author notes:** joint last authors (, and) (David Presby, Dennis Bontempi, Sven Bergmann). **Corresponding author** | Sacha Bors, Department of Computational Biology; Genopode, office 2025.1 - CH1015 Lausanne, Switzerland.

## Abstract

Deep learning models based on Vision Transformers (ViTs) have shown strong performance in retinal fundus imaging, but their interpretability remains poorly understood. In particular, attention-based attribution methods are widely used to explain ViT predictions, despite limited evaluation of their faithfulness and biological relevance in medical imaging. Here, we systematically benchmark four attention-based interpretability methods for RETFound, a retinal ViT-based foundation model, that we previously fine-tuned to predict 17 retinal vascular phenotypes from UK Biobank fundus images^1^. We compare raw attention, attention rollout, gradient-weighted attention rollout, and Chefer’s hybrid relevance-based method using both qualitative visualisation and quantitative evaluation frameworks. To assess attribution faithfulness, we perform perturbation-based deletion and insertion experiments, quantifying changes in model predictions as highly attended image regions are progressively removed or restored. To evaluate biological specificity, we run structure-aware analyses combining attribution maps with vessel segmentation and artery-vein labels through the Relative ratio of Attention Intensity (RAI) metric. Across models, attribution maps differed substantially depending on the selected interpretability method, highlighting the need for rigorous quantitative evaluation. Among the evaluated approaches, gradient-weighted attention rollout consistently achieved the strongest perturbation performance and produced attribution maps most closely aligned with the anatomical definition of the predicted retinal traits. Furthermore, vessel-type specific models systematically concentrate attention on the corresponding vascular structures despite being trained using only a single scalar value per image as supervision. These findings demonstrate that attention-based attribution methods capture biologically meaningful vascular representations, while also revealing method-dependent variability in attribution behaviour. This work provides a quantitative framework for evaluating interpretability methods in medical imaging with annotated segmentation and contributes toward more transparent and biologically grounded medical AI systems.

## Introduction

The retinal vasculature provides a unique and accessible window into the human vascular system. Unlike most vascular beds, retinal vessels can be visualised non-invasively using colour fundus photography, allowing direct observation of vasculature at a large scale. Numerous studies have demonstrated that retinal vascular features are associated not only with ocular diseases such as diabetic retinopathy, macular degeneration, and glaucoma^2^, but also with systemic conditions, including hypertension^3,4^, cardiovascular^5–7^ and cerebrovascular disease^8^, diabetes^9^, and overall mortality^10,11^. As a result, retinal fundus images have become an increasingly valuable modality for population-level studies of vascular health.

Large biobank cohorts have further amplified this potential. In particular, the UK Biobank has collected retinal fundus images for over 130,000 eyes from approximately 72,000 participants, along with phenotypic, clinical, and genetic data. This combination enables systematic investigation of how retinal vascular morphology relates to disease risk and genetic architecture at the population scale. However, extracting biologically meaningful information from such large image collections requires automated and reproducible analysis pipelines. In this context, Ortín Vela et al.^12^ developed a fully automated framework to extract and analyse 17 retinal vascular phenotypes from fundus images, including vessel diameters, diameter variability, tortuosity, vascular density, bifurcation counts, temporal angles, and central retinal equivalents. When applied to the UK Biobank, this framework enabled the estimation of SNP-based heritability for these features and revealed numerous genetic associations and disease correlations.

Despite their interpretability and clinical relevance, measured image features are subject to several inherent limitations. For instance, they capture only a predefined subset of image information and rely on heuristic choices made during segmentation, skeletonisation, and feature computation, which may not generalise uniformly across imaging conditions or populations. In parallel, deep learning (DL) approaches have emerged as powerful alternatives for analysing retinal images. Convolutional neural networks and, more recently, Vision Transformers^13,14^ have been successfully applied to tasks ranging from disease classification to risk prediction^15^. Unlike classical pipelines, these models learn complex, high-dimensional representations directly from images, enabling them to capture subtle and distributed patterns that are difficult to formalise manually, in a way that is often more robust to noise and image variability.

Foundation models represent a recent and particularly influential development in this direction. Pretrained using self-supervised objectives on large image collections, such as masked image reconstruction, these models learn general representations that can be fine-tuned for diverse downstream tasks in a second training step. RETFound is a transformer-based foundation model specifically developed for retinal imaging and pretrained on large-scale fundus datasets using a masked autoencoder framework (Zhou et al.^16^). Beyeler et al.^1^ leveraged RETFound to systematically compare deep learning-derived representations with classical retinal vascular phenotypes in the UK Biobank. By fine-tuning RETFound to predict the 17 tangible vascular features defined by Ortín Vela et al.^12^, we showed that some features, such as vascular density, can be predicted with high accuracy, whereas more complex features, such as tortuosity, are only partially captured. Importantly, genome-wide association analyses revealed that RETFound’s latent space variables carry genetic signals that are distinct from those of classical vascular phenotypes, suggesting that deep learning models might encode complementary biological information.

While these results highlight the promise of deep learning for retinal phenotyping, they also raise important questions regarding interpretability. Deep neural networks, and foundation models in particular, are often criticised as “black boxes”, providing accurate predictions without clear explanations of which image features drive those predictions. This lack of transparency limits clinical trust, hinders biological interpretation, and complicates rigorous model validation and assessment. In response, an expanding literature has proposed explainability methods, such as saliency and attention-based visualisation techniques^17–19^, which aim to identify image regions that contribute most strongly to a model’s output.

In retinal imaging, attention and saliency maps have been used to assess whether models focus on clinically relevant structures^16,20–24^ such as blood vessels, the optic disc, or the macula. For instance, Trofimova et al.^25^ employed attention-based visualisations to interpret RETFound models fine-tuned to predict chronological age from fundus images, using these analyses to characterise sex-specific differences in model focus and biological associations, including a stronger vascular signal in females. Such studies illustrate how explainability analyses can support biological interpretation of deep learning models. However, it has also been shown that attribution methods can be sensitive to methodological choices, potentially unstable across implementations, and that visually plausible heatmaps do not necessarily reflect the true internal mechanisms of a model^26^.

Despite the increasing use of explainability methods in ophthalmic deep learning, systematic benchmarking of their faithfulness and biological specificity remains limited. In particular, little is known about how different attention-based interpretability methods behave when applied to Vision Transformer architectures fine-tuned on well-defined, biologically grounded retinal phenotypes. Moreover, many studies rely primarily on qualitative assessment of attribution maps, often illustrated through a small number of selected examples, and quantitative evaluation strategies that extend beyond visual inspection are still rarely applied.

This work aims to address these limitations in the context of retinal vascular phenotyping from fundus images. Building on the set of 17 features defined by Ortín Vela et al.^12^ and the fine-tuned RETFound models released by Beyeler et al.^1^, we conduct a systematic evaluation of attention-based interpretability methods in retinal fundus imaging. Using the same UK Biobank dataset and phenotype-specific models, we investigate how different attribution strategies highlight image regions, how faithfully they reflect model reliance on image content, and whether they capture biologically meaningful vascular structure.

Specifically, this work benchmarks multiple attention-based interpretability methods for RETFound fine-tuned models, including raw attention, attention rollout, gradient-weighted attention rollout, and a hybrid relevance-based approach^27^. Beyond qualitative visualisation, we introduce quantitative evaluation frameworks based on perturbation experiments and structure-specific attention metrics, enabling direct comparison of attribution methods across

17 distinct vascular phenotypes. By integrating vessel segmentation masks, artery-vein labels, and control models with randomised parameters, these analyses aim to disentangle learned biological signals from patterns arising from methodological biases and to provide evidence that the resulting attribution maps reflect meaningful aspects of the models’ learned representations.

The goal of this research is not only to assess which interpretability methods perform best under specific conditions, but also to clarify what deep learning models trained on retinal images actually learn about vascular biology. Ultimately, it aims to contribute to the development of more transparent, trustworthy, and biologically interpretable AI models for retinal imaging and vascular health research.

## Results

### Qualitative Evaluation

We systematically computed attribution maps for all 17 phenotype-specific models using four interpretability methods: *raw attention*, *attention rollout*, *gradient-weighted attention rollout*, and *Chefer’s hybrid method*^27^ (see the Methods Section). We observed a large variation in attribution patterns across the 17 predictive targets, consistent with models learning distinct spatial representations. For arterial and venous vascular density (**Figure 1a,b**), attribution maps predominantly highlighted vessel structures, but with notable differences between vessel types. In particular, the venous vascular density model showed less focus on vessels close to the macular region. Models predicting temporal angles showed a localised focus on regions near the optic disc (**Figure 1c**), closely aligning with the definition of these traits^12^, which are computed from vessel endpoints detected at a fixed radius around the optic disc. We also observed that the model trained to predict bifurcation count showed distributed attribution (**Figure 1d**) across both the optic disc and distal vasculature, with emphasis on higher-order vessel branches.

**Figure 1.**
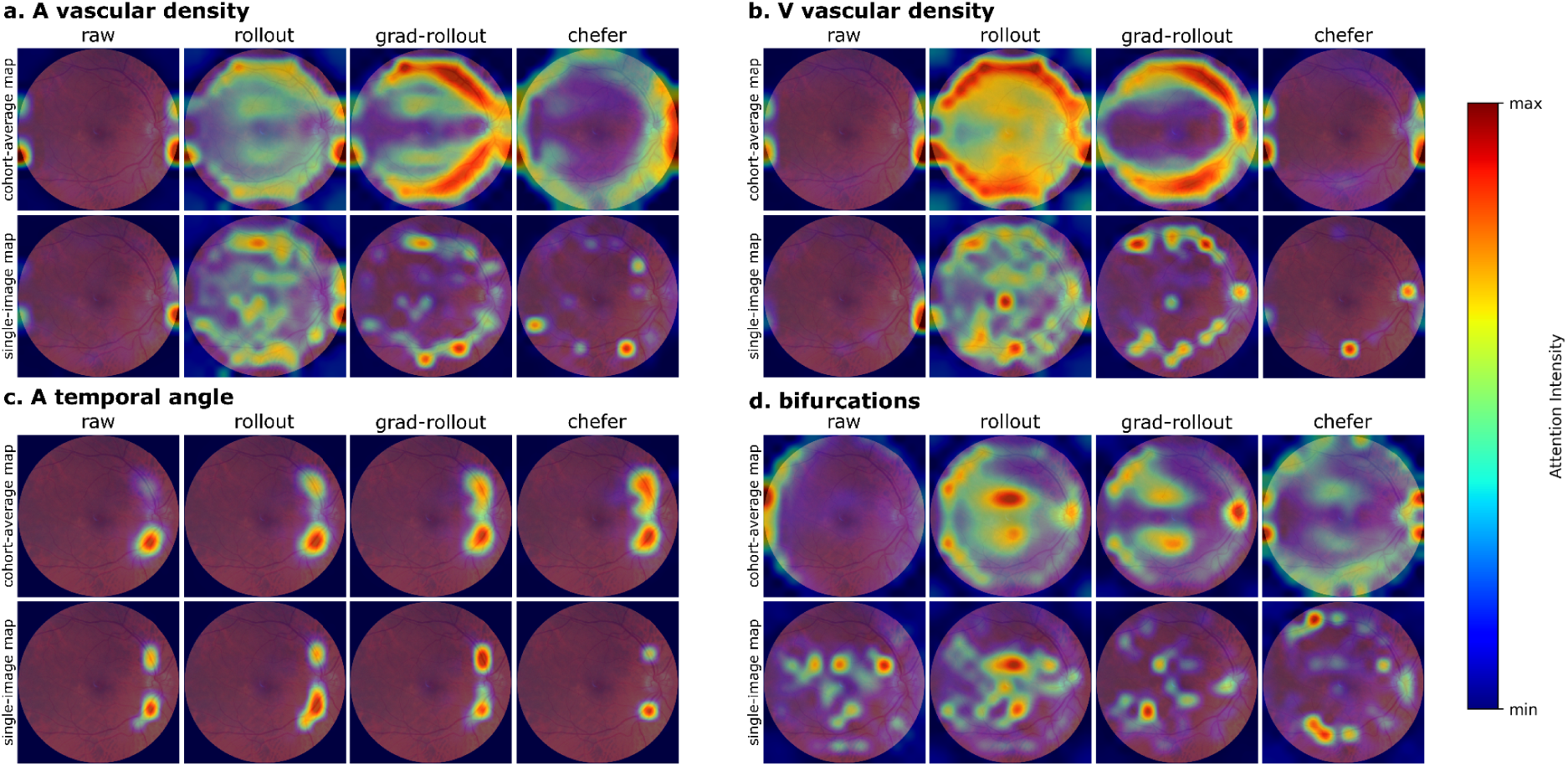
Qualitative comparison of attribution maps across phenotypes and interpretability methods. Attribution maps are shown for four models: **(a)** arterial vascular density, **(b)** venous vascular density, **(c)** arterial temporal angle, and **(d)** bifurcation count. For each phenotype, attribution maps obtained using four different methods (raw attention, attention rollout, gradient-weighted attention rollout, and Chefer’s hybrid method) are displayed. The top row in each panel shows cohort-averaged attribution maps, while the bottom row shows attribution maps for a representative individual image, all overlaid on the same reference fundus image.

**Figure 2.**
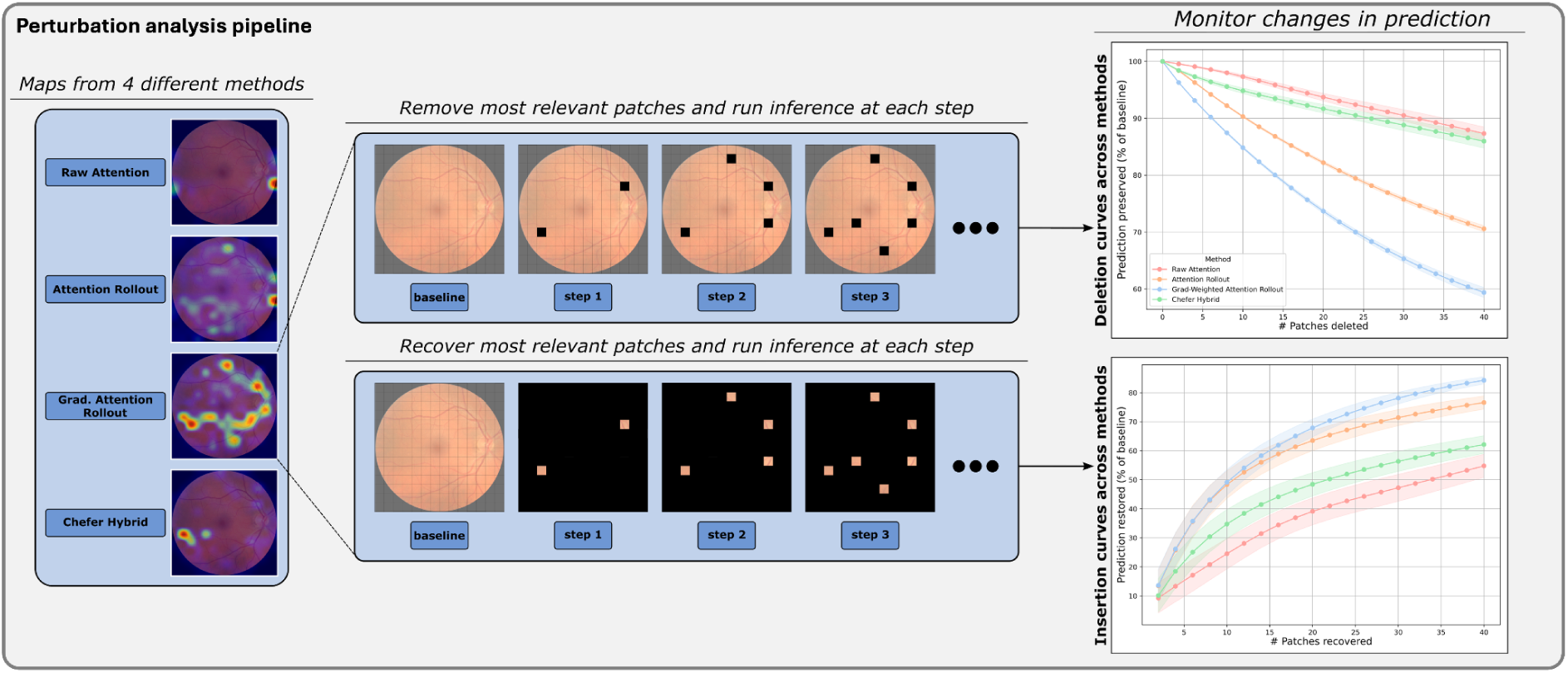
Perturbation analysis pipeline. Attribution maps generated by four interpretability methods (raw attention, attention rollout, gradient-weighted attention rollout, and Chefer’s hybrid method) were used to rank image patches according to their attribution score. In deletion experiments (top), the most attended patches were progressively removed from the image, whereas in insertion experiments (bottom), they were progressively restored starting from a fully masked image. Prediction curves obtained across perturbation steps were then used to quantify how strongly model predictions depend on the regions highlighted by each attribution method.

Beyond these phenotype-specific patterns, attribution methods did not consistently agree, neither at the individual image level nor when averaged across the cohort. Combining cohort-level and single-image visualisations therefore provides complementary insight into global model behaviour as well as into how attribution manifests in concrete examples. In particular, cohort-level attribution maps are informative in this context, since colour fundus images are spatially well-aligned across individuals (only right eyes), with the fovea typically centred and the optic disc on the nasal side (on the right). The lack of systematic agreement across methods motivates the need for quantitative evaluation to assess which attribution strategies most faithfully reflect the model’s learned representations.

### Perturbation-Based Evaluation

We evaluated how faithfully the different attribution methods identify image regions that drive model predictions using deletion and insertion perturbation experiments^28–31^ across all 17 fine-tuned models. For each model and attribution method, predictions were systematically perturbed by progressively removing or reintroducing the most highly attended regions, allowing us to quantify how strongly the model’s output depended on the regions highlighted by each attribution technique (Figure 3). We summarised the effect of perturbation using the area above the deletion curve and the area under the insertion curve, providing a single scalar measure of attribution faithfulness per model and method (Figure 4).

Across models, we did not observe a single attribution method that uniformly outperformed all others in every setting. Instead, the relative performance of attribution methods varied depending on the predicted phenotype, indicating that attribution faithfulness is model- and task-dependent. When considering performance aggregated across all 17 models, attribution methods were ranked according to their deletion scores for each phenotype. Gradient-weighted attention rollout achieved the best overall performance, with an average rank of 1.35 out of 4 across models. This method significantly outperformed all other attribution methods for 11 out of 17 models (SEM across folds), indicating a more reliable identification of regions impacting the model’s predictions.

**Figure 3.**
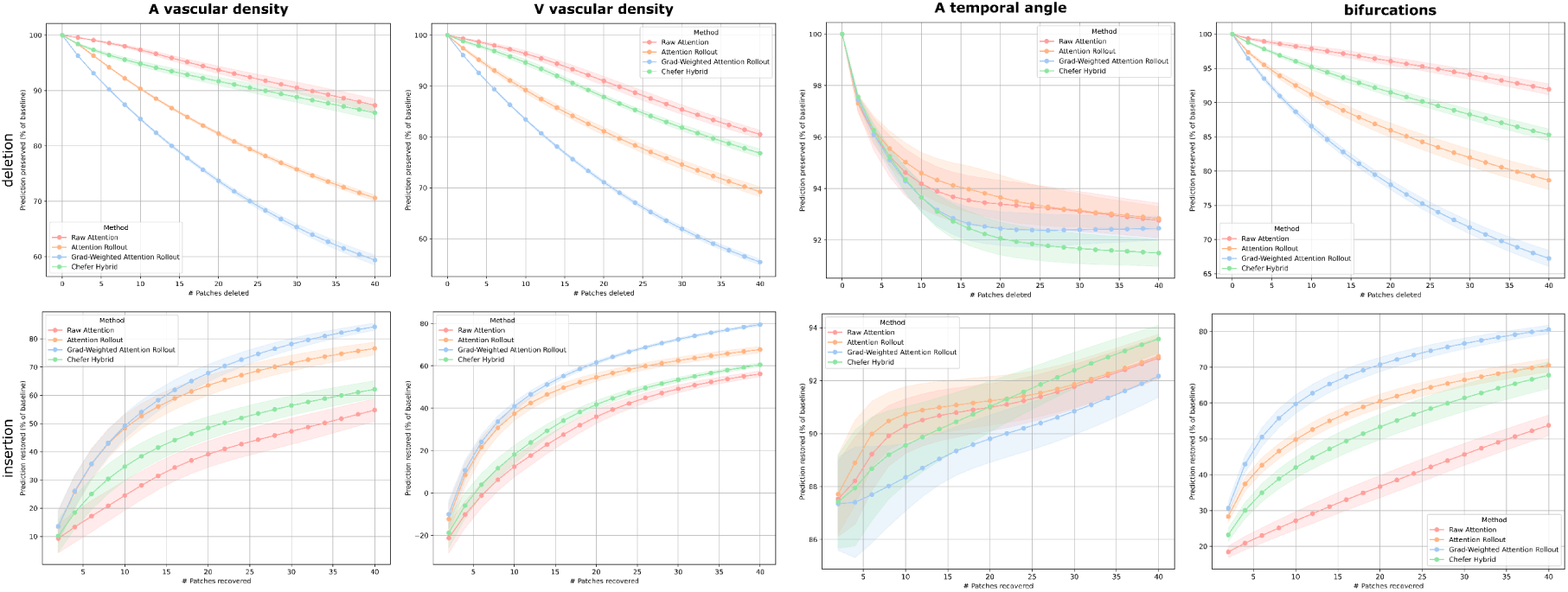
Perturbation curves across attribution methods and predicted traits. For each subplot, we show prediction preservation or restoration (mean ± SEM across folds) as a function of the number of perturbed patches (2-patch steps, up to 40 patches, i.e. ∼20% of the image) - expressed in percentage of baseline prediction. Top: deletion curves, prediction drop when iteratively removing the most attended patches. Bottom: insertion curves, prediction recovery when reintroducing patches.

**Figure 4.**
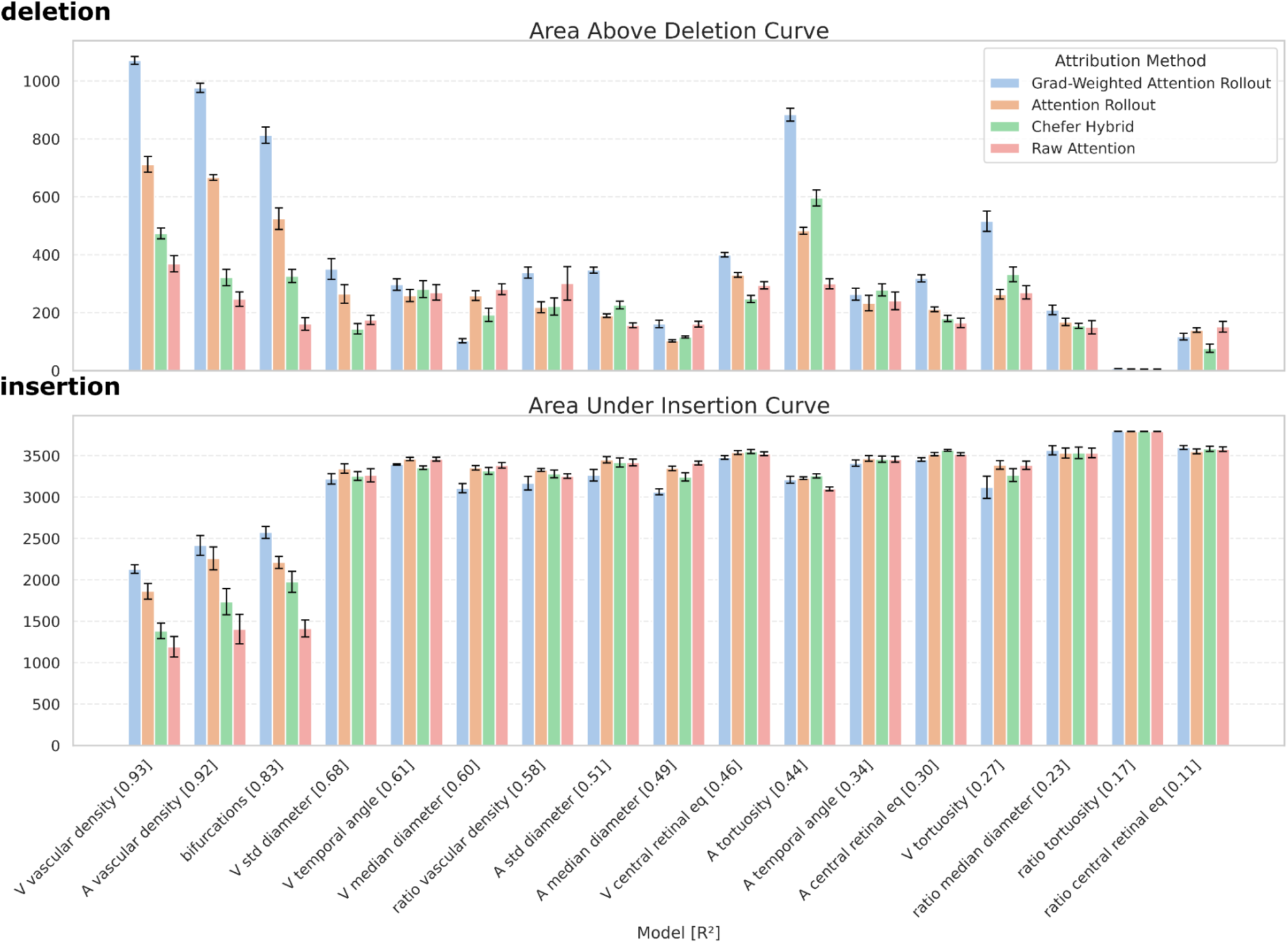
Perturbation-based evaluation of attribution methods across models. Bar plots show the mean perturbation score (±SEM across folds) for each attribution method (Grad-Weighted Attention Rollout, Attention Rollout, Chefer Hybrid, and Raw Attention), computed over 17 fine-tuned RETFound models. (top) Deletion: area above the deletion curve obtained by iteratively removing the most attended patches. (bottom) Insertion: area under the insertion curve obtained by iteratively reintroducing the most attended patches. Models are ordered from left to right by decreasing predictive performance (R^2^).

We further observed that the three models with the highest predictive performance (top R^2^: 0.93 - V vascular density, 0.92 - A vascular density, and 0.83 - bifurcations, respectively) exhibited identical rankings both in deletion and insertion experiments. For these models, gradient-weighted attention rollout produced the steepest early prediction drop in deletion and the fastest recovery in insertion, followed by attention rollout, Chefer’s hybrid method, and raw attention. This consistent ordering across both perturbation tests suggests that, for well-learned phenotypes, attribution faithfulness becomes more stable and less dependent on the specific perturbation test. These top-performing models also showed larger overall perturbation effects across all attribution methods, with higher areas above the deletion curve, indicating that stronger predictive models are more sensitive to perturbations of informative image regions.

In contrast, for models with lower predictive performance (R^2^ in the range from 0.68 to 0.11), differences between attribution methods became less pronounced, particularly in the insertion experiments. For these models, insertion curves largely overlapped across methods, and no statistically significant differences were observed.

### Vascular Structure Analysis

To further characterise the biological specificity of the attribution maps, and to assess whether the models capture meaningful concepts beyond their original training targets, we quantified how attention was distributed across vascular structures using the *Relative ratio of Attention Intensity* (RAI) metric^21^. This analysis uses attribution maps together with vessel segmentation masks^32^ to compare the average attribution assigned to vessel pixels and to specific vessel types. RAI provides a direct, quantitative ratio describing how strongly a model concentrates its attention on arteries or veins relative to complementary regions of the image. By analysing the distribution of RAI values across images, models, and attribution methods, we assessed whether attention maps consistently reflect the vascular nature of the predicted traits and whether vessel-type specific models highlight their corresponding vascular structures.

Across all 17 models and attribution methods, we consistently observed RAI mean values greater than 1 for vasculature as a whole, indicating that attribution maps were systematically concentrated on vessel regions compared to the non-vascular background (Figure 5a). As a consequence, RAI mean values for non-vessel regions were consistently below 1. This pattern was observed irrespective of predictive performance and attribution method, suggesting that all models base their predictions mainly on vascular information.

**Figure 5.**
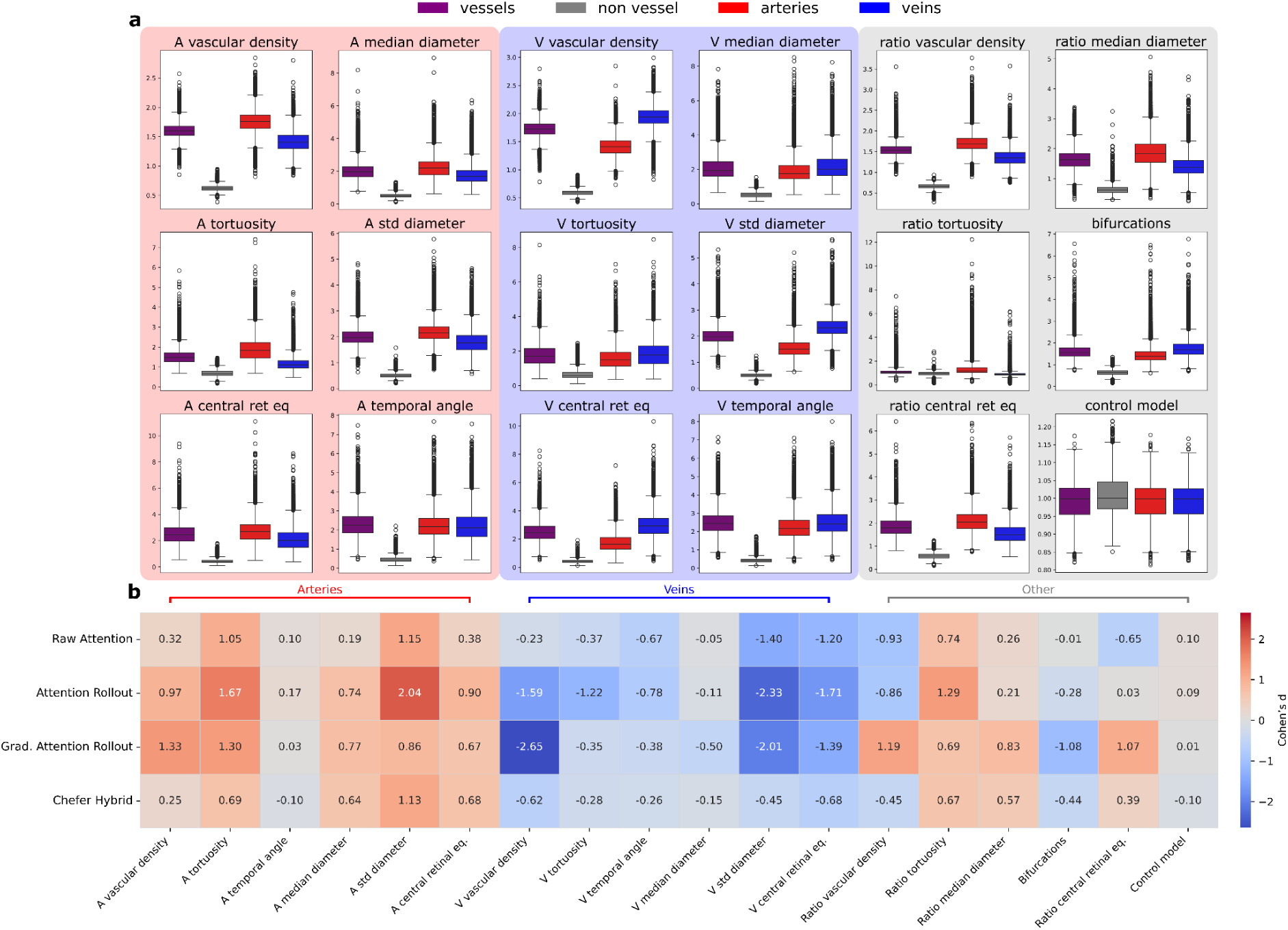
Relative ratio of Attention Intensity (RAI) across vascular structures and vessel types. **(a)** Boxplots of RAI computed for four anatomical structures (vessels, non-vessel regions, arteries, and veins) across all 17 fine-tuned models using gradient-weighted attention rollout maps. Each panel corresponds to a separate model. RAI distributions obtained from the control model are shown in the bottom right panel. **(b)** Cohen’s d effect sizes quantifying the difference in RAI between arteries and veins across all models and attribution methods. Each cell shows the standardised mean difference in RAI (arteries − veins), computed across all images. Positive values indicate higher relative attention on arteries compared with veins, and vice versa for negative values. Models are grouped by feature type (arterial, venous, and other) and compared across four interpretability methods: raw attention, attention rollout, gradient-weighted attention rollout, and Chefer’s hybrid method. The control model corresponds to a randomly initialised RETFound model.

Beyond global vasculature, we examined whether attribution maps captured vessel-type specificity. For phenotypes defined on a single vessel type, such as artery- or vein-specific traits, models consistently exhibited higher RAI values for the corresponding vessel type than for the other. When quantified using Cohen’s d, this separation was large and consistently oriented in the expected direction for vessel-specific traits (Figure 5b). The largest effect was observed for venous vascular density, with Cohen’s d = −2.65, while arterial vascular density also showed a large effect (d = 1.33). Similarly, very large effect sizes were found for the standard deviation of diameter, with Cohen’s d values reaching 2.04 for arteries and −2.33 for veins. In contrast, the arterial temporal angle model was the only vessel-type specific model to show consistently small effect sizes (d ≤ 0.17), which may reflect the localised nature of the corresponding attribution maps and the fact that patches adjacent to the optic disc, where these angles are defined, contain both arteries and veins.

Together, these results demonstrate that vessel-type specific models reliably learn to distinguish arteries from veins, even though training supervision is provided only through a single scalar phenotype encoding complex vascular information.

For phenotypes that by definition combine information from both vessel types, such as bifurcation counts or ratio-based features, no uniform pattern emerged across traits. Instead, RAI distributions and corresponding Cohen’s d values varied across models and attribution methods. For the ratio of tortuosity and the ratio of median diameter, all four attribution methods consistently indicated a predominance of attention on arteries. In contrast, for the bifurcation count, the majority of attribution methods showed a stronger allocation of attention to veins. For the ratio of vascular density and the ratio of CRE, attribution methods did not consistently agree on the vascular structure receiving greater attention.

Finally, applying the same RAI analysis to the control model (i.e., initialised with randomised parameters) revealed no significant separation between artery and vein RAI distributions. Cohen’s d values for the control model were close to zero across attribution methods, indicating a failure to differentiate between vessel types in the absence of learned representations. Together with the results observed from the trained models, this provides strong evidence that vessel-type specificity in attribution maps arises from learned structure, rather than from image artefacts, biases of the attribution methods themselves, or of the RAI measurement.

## Discussion

In this study, we systematically evaluated attention-based interpretability methods for Vision Transformers in retinal fundus imaging, and showed that variability between attribution maps remains substantial, both at the level of individual images and cohort-averaged maps. Except for highly localised tasks such as temporal angle prediction, different attribution methods often highlighted markedly different retinal regions despite being applied to the same model and prediction target. Together, these findings highlight the need for rigorous quantitative evaluation when using attribution maps to derive biological or clinical interpretations from deep learning models.

Among the evaluated approaches, gradient-weighted attention rollout consistently emerged as the most faithful attribution method overall. Across the perturbation analyses, this method achieved the best average ranking, indicating that the highlighted regions were more strongly associated with the model’s predictions. Qualitatively, gradient-weighted attention rollout also generated attribution maps that most closely matched the expected spatial definition of the underlying retinal phenotypes. For vascular density prediction, attribution maps predominantly followed vascular structures, whereas for temporal angle and central retinal equivalent prediction, attention concentrated around optic-disc-adjacent regions corresponding to the geometric definition of the traits. These findings suggest that combining attention scores with attention gradients across transformer blocks improves the identification of image regions that genuinely contribute to the model’s output, and more faithfully captures the effective flow of information through the network during inference.

Our perturbation analyses revealed that the interpretability and stability of attribution methods strongly depend both on the predictive performance of the underlying model and on the nature of the predicted phenotype. The three models with the highest R^2^ exhibited a notable consistency between deletion and insertion experiments, with all attribution methods following the same ranking. In contrast, lower-performing models showed substantially smaller differentiation between methods, particularly during insertion experiments, where perturbation curves frequently overlapped. One possible explanation is that insertion experiments are inherently more challenging than deletion experiments, as predictions must be recovered from only a small fraction of the original image. As a result, insertion curves may provide less discriminative power between attribution methods, particularly for models with weaker predictive performance. Furthermore, when baseline predictions are already noisy, perturbed predictions may remain relatively close to the original output. This may result in inflated performance estimates, providing less discriminative power between attribution methods for these specific cases (i.e., low R^2^). In our perturbation experiments, the models’ behaviour also appeared to depend on the spatial nature of the predicted phenotype itself. Predictions for localised phenotypes recovered rapidly as the most highly attended regions were reintroduced, whereas predictions for more spatially distributed phenotypes required substantially larger fractions of the image to achieve comparable recovery. Such differences were consistent with the anatomical characteristics of the target phenotypes: if localised traits, such as temporal angles, can plausibly be recovered from only a few highly informative patches, other ones, such as median vessel diameter, are distributed across much larger portions of the retinal vasculature and therefore might require more extensive image information for accurate prediction recovery. This suggests that perturbation-based evaluation should not be interpreted independently of both model performance and the spatial complexity of the target phenotype.

Overall, our vascular structure analyses demonstrate that fine-tuned RETFound models, despite being trained exclusively to predict a single image-level vascular phenotype, implicitly learn biologically meaningful vessel-type representations. Across all attribution methods, models consistently concentrated attention on vascular regions rather than background tissue, and vessel-type specific models systematically focused more strongly on the corresponding vascular structures. Models trained on arterial phenotypes predominantly attended to arteries, whereas models trained on venous phenotypes predominantly attended to veins, producing large Cohen’s d effect sizes, whose signs align with our expectations, across artery-vein comparisons. These vessel-type differentiation patterns disappeared in the randomly initialised control model, indicating that they emerged from learned representations rather than from image properties, architecture characteristics, or biases of the attribution methods themselves. These findings therefore suggest that, while learning to predict complex retinal phenotypes such as tortuosity or vascular density, the models simultaneously acquire secondary biological concepts, including vessel identity and vascular organisation. More broadly, this provides an important proof of concept for applying attribution analyses to models trained on more complex clinical outcomes, such as disease prediction or systemic risk-factor estimation, where the relevant biological mechanisms may not be known a priori. In such settings, attention-based analyses could help identify anatomical regions or vascular structures that contribute most strongly to prediction, potentially increasing confidence in model behaviour and generating hypotheses regarding previously unknown biological associations. Similar approaches have already been explored in retinal disease prediction studies^22^, where saliency analyses suggested that small vessels and capillaries may contribute strongly to Alzheimer’s disease classification from retinal vasculature images, which aligns with the known degeneration of small vascular structures associated with neurodegenerative diseases.

Several limitations of this study should nevertheless be acknowledged. First, our benchmark focuses on attribution faithfulness and biological specificity, but does not assess the robustness of attribution maps to image perturbations or acquisition variability. Future work could therefore investigate the stability of attention maps under changes in image quality, brightness, geometric transformations, or other clinically relevant perturbations. Second, our analyses are restricted to models trained to predict retinal vascular phenotypes, whose anatomical correlates are relatively well understood. Whether the same conclusions extend to disease prediction or systemic risk-factor estimation remains to be determined. Finally, we focused exclusively on attention-based interpretability methods. Comparing these approaches with other families of explainability techniques may help establish a more comprehensive framework for interpreting retinal foundation models.

## Methods

### Model Training & Setup

For all predictive analyses, we used RETFound, a transformer-based foundation model for retinal imaging introduced by Zhou et al.^16^. RETFound is built upon a ViT-L/16 encoder^14^ trained using a masked autoencoding framework, in which large portions of the input image are masked, and the model is optimised to reconstruct the missing patches. This self-supervised procedure enables the encoder to learn generic retinal representations from large-scale unlabelled datasets.

The target phenotypes used in this study correspond to the 17 retinal vascular image features previously defined and measured by Ortín Vela et al.^12^. These features, including vessel diameters, temporal angles, tortuosity, vascular density, and bifurcation count, were extracted using the fully automated pipeline described in the publication and were used without modification.

The fine-tuned RETFound models employed in this study were obtained directly from Beyeler et al.^1^, who fine-tuned RETFound to predict each of the 17 features by attaching a multilayer perceptron head to the pretrained encoder. The authors fine-tuned both the encoder and the prediction head simultaneously. This training followed the hyperparameter configuration established in the original RETFound manuscript: 50 epochs with a batch size of 16, an initial 10-epoch linear learning-rate warm-up from 0 to 5 × 10^-4^, followed by a smooth decay to 1 × 10^-6^, a dropout rate of 0.2, and image inputs resized to 224×224 pixels. For each image feature, models were trained using 5-fold cross-validation, with data split into 60%, 20%, and 20% training, validation, and test subsets, respectively, in each fold. In the present work, all interpretability, attribution, and downstream analyses were performed on the same fine-tuned models as released by the authors^1^, without further training or modification.

### Models Architecture

All fine-tuned models are based on the Vision Transformer Large architecture with 16×16 patching (ViT-L/16). With this architecture, images are decomposed into 196 patches, which are projected into a 1024-dimensional embedding space via a convolutional patch-embedding layer. A learnable class (CLS) token is added, resulting in a sequence of 197 tokens.

The transformer encoder consists of 24 identical blocks. Each block applies multi-head self-attention with 16 attention heads, followed by a multilayer perceptron (MLP) with an expansion of the embedding dimension (1024→4096→1024) and GELU activations. Layer normalisation is applied before both attention and MLP layers.

For prediction, token representations are aggregated using global average pooling over patch tokens, rather than relying solely on the CLS token. The pooled representation is passed through a final normalisation layer and a linear regression head, producing a single scalar output. Each model contains approximately 303 million trainable parameters and shares the same architecture.

### Attention Extraction

To analyse and compare attention-based interpretability methods, we extracted attention maps for all of our fine-tuned models. Specifically, we implemented a pipeline supporting four interpretability modes: (1) **Raw Attention**, (2) **Attention Rollout**^17^, (3) **Gradient-weighted Attention Rollout**, and (4) **Chefer’s hybrid method**^27^. Each mode yields a 14×14 spatial attention map per image that corresponds to the CLS-to-patch attention, which captures how much each image patch attends to the classification token. The extracted maps reflect the spatial distribution of the information that directs the model’s focus during inference.

#### Raw Attention

We extracted the attention matrix from the final transformer block (block 24), directly after the softmax activation. This final layer encodes the features with the highest level of abstraction. In order to weight attention heads according to their influence on the model prediction, instead of uniformly averaging them, we applied a head sensitivity analysis (described in a following section) to derive the weights *w_h_*, which reflect the contribution of each attention head to the output prediction.

The attention matrix from head *h* at layer *l* is computed as:

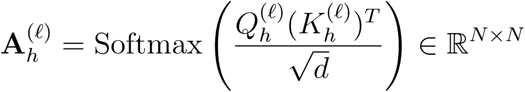

where 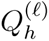 and 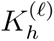 are the query and key matrices, *N* = 197 is the number of tokens (1 CLS + 196 patches), and *d* = 64 is the attention head dimension. The final attention matrix is a weighted average across heads in the last layer (i.e., *H* = 16 and *L* = 24):

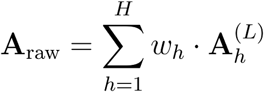

We then extracted the CLS-to-patch vector^33^ **A**_raw_[0,1:] (first row, excluding the first column), corresponding to the attention from the classification token to each image patch. The CLS-to-CLS attention value was excluded because it has no spatial correspondence in the input image. The remaining 196 values were reshaped into a 14×14 grid.

#### Attention Rollout

This method, introduced by Abnar & Zuidema^17^, recursively combines attention from all transformer layers. At each layer, we average attention heads and add the identity matrix **I** to account for residual connections:

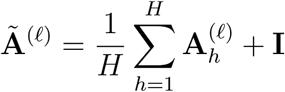

The final rollout attention is obtained by recursively multiplying these matrices:

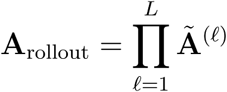

We again extract the CLS-to-patch attention vector and reshape it into a 14×14 grid.

#### Gradient-weighted Attention Rollout

Inspired by Chefer et al.^27^, this method modulates attention matrices using their gradients with respect to the output. For each head *h* and layer *l*, we compute:

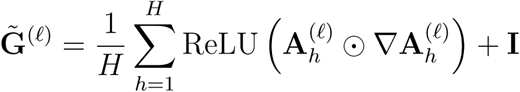

where ReLU (·) denotes the rectified linear unit activation function. These modified matrices are then aggregated through rollout:

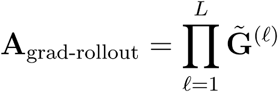

#### Chefer’s Hybrid Method

We adapted and used the code from Chefer et al.^27^ to compute relevance scores via Layer-wise Relevance Propagation^34^ (LRP). This method propagates relevance scores from the output back to the input through the whole transformer architecture, with customised rules for each type of layer. This allows us to attribute a proportion of the final output, layer by layer, to every single node of the model. Originally defined for CNNs, the authors^27^ further implemented LRP rules for ViTs, including attention layers and normalisation layers. Following Chefer’s hybrid method, we placed a hook on relevance scores 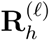 backpropagated to attention layers, multiplied them with attention gradients, and averaged across heads as follows:

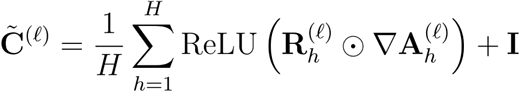

Final attribution maps are obtained through rollout:

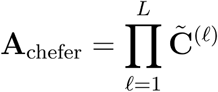

### Visualisation of Attribution Maps

For all interpretability methods, we visualised the resulting 14×14 attribution maps using a common procedure. Each attribution map encodes patch-level importance (one value per 16×16 input region). For *raw attention*, which produces 16 head-specific maps, we first aggregated heads using the per-image head-importance weights derived from the head-sensitivity analysis. The other three methods, *attention rollout*, *gradient*-*weighted rollout*, and *Chefer’s hybrid method*, already return a single map per image and therefore require no head aggregation.

Since attribution maps are primarily interpreted through their relative rather than absolute values, we normalised each attribution map by its own minimum and maximum value before visualisation. This scaling preserves the relative spatial distribution of attention within each map while making individual examples visually comparable. The 14×14 maps were then upsampled to full resolution (224×224) using cubic interpolation, and overlaid on the corresponding reference fundus image.

We visualised attribution maps at two levels:

1. **Individual maps**, allowing direct inspection of the spatial regions attended by the model on a specific fundus image.
2. **Cohort aggregated maps**, obtained by averaging the normalised attribution maps across all images. Since fundus images are spatially well-aligned across individuals (fovea-centred images with optic disc on the right), this averaging reveals stable, population-level patterns in the model’s focus during inference, without relying on single examples.

### Head-sensitivity Analysis

Raw attention maps in Vision Transformers reflect how information is distributed across attention heads, but they are not conditioned on the model output and therefore do not directly indicate which heads are functionally important for a particular prediction^35^. A common practice is to average the attention heads in the final block uniformly, but this implicitly assumes that all heads contribute equally to the prediction, an assumption that is unlikely to hold in practice. To introduce output dependence into raw attention aggregation, we performed a head-sensitivity analysis that quantifies, for each image, how much the model’s prediction relies on each head in the last transformer layer (24th block).

For each trained model, and for each of the 16 heads in the last attention block, we created a perturbed version of the model in which that head’s attention weights were zeroed out during inference. This removes the contribution of a specific head while leaving the rest of the model unchanged. We then ran inference on the full cohort and compared the perturbed prediction to the baseline prediction from the unmodified model for every image. For the image *i* and head *h*, we computed the absolute difference

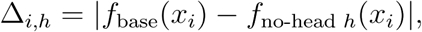

which measures how strongly the prediction for that specific image depends on head *h*.

We obtained, for each image, a 16-dimensional vector of head-specific deltas. We then used the normalised squared deltas to compute head-importance weights:

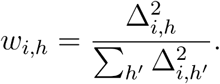

These weights increase the importance of attention heads whose removal produces large prediction changes and vary from one image to another, while reducing that of heads that don’t meaningfully influence the output.

To obtain a raw attention map, we computed a weighted average of the 16 head-specific attention maps using these per-image weights, instead of a linear average.

### Perturbation-Based Evaluation

To assess the faithfulness of the extracted attention maps, i.e. how well they highlight regions that genuinely influence model predictions, we implemented a perturbation-based evaluation inspired by standard interpretability benchmarks^28–31^. For each attention method, we conducted prediction perturbation experiments that measure the impact of the masking or revealing of patches considered important by the attribution map.

For each image in the cohort and each attribution method, we performed two types of perturbations: deletion and insertion. In the deletion experiment, we first ranked the 196 patches of the input image according to their attribution score. Starting from the original image, we progressively masked the top-k most attended patches (in steps of 2 patches, up to 40, masking ∼20% of the original image) by setting their pixel values to zero, and re-ran model inference at each step. We recorded the absolute prediction change compared to the unperturbed image (baseline), expressing it as a percentage of the baseline prediction. This yielded a prediction preservation curve that started at 100% (no perturbation) and decreased as highly attended patches were deleted. A steeper early decline indicates that the method successfully prioritised patches essential to the model’s prediction, providing evidence of a stronger correspondence between the attribution map and the model’s underlying decision process.

Conversely, in the insertion experiment, we reversed the process. Starting from a fully masked image, we progressively restored the top-k most attended patches in steps of two (recovering up to 40 patches, i.e. ∼20% of the image). This produced a prediction restoration curve that increased from a low initial value toward the baseline prediction. Here, a steeper early increase similarly suggests a more faithful attribution method.

While the deletion experiment measures how sensitive the model’s prediction is to the removal of the most attended regions, the insertion experiment instead quantifies how quickly the model can recover its prediction when given access to those same regions.

For each model and attribution method, we quantified the effect of perturbation by first computing, for every image in the cohort, the absolute change in prediction relative to its baseline value. These per-image differences were then averaged across the cohort to produce a single prediction-preservation curve per fold. By repeating this procedure for each of the five cross-validation folds, we obtained five curves per attribution method. These were then aggregated by computing the mean curve across folds, and we reported the standard error of the mean (SEM) to reflect variability. Finally, to summarise overall performance with a single scalar metric, we computed the area above the curve (for deletion experiments) and the area under the curve (for insertion experiments), with larger values indicating stronger predictive degradation or recovery when perturbing regions highlighted by the attribution method.

### Vascular Structure Analysis

To quantify the extent to which models focus on specific vascular structures during inference, we computed the *Relative ratio of Attention Intensity* (RAI), a metric adapted from Lee et al.^21^. For each image, we overlaid the attribution map produced by a given model and interpretability method with the corresponding vessel segmentation mask, which labels each pixel as belonging to arteries, veins, overlap of both, or background (non-vessel). The vessel masks used in this analysis were obtained from a state-of-the-art automated segmentation pipeline^32^. To improve vascular connectivity and reduce artefacts, we applied a deep-learning-based vessel reconstruction step using an iterative post-segmentation process^36^. The attribution maps were normalised to sum to one across the entire image.

Given a binary mask for a target structure *S*, we computed the RAI as the ratio between the average attention score within the structure *S* and the average attention score outside of it:

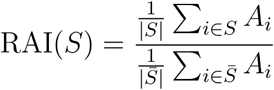

where *A_i_* is the attention score at pixel *i*, *S* denotes the set of pixels in the target structure, and *S̄* its complement. An RAI value of ℓ indicates that, on average, attention is ℓ times more concentrated per pixel inside the structure than outside. This procedure yields one RAI value per image and per structure, allowing us to compare the resulting distributions across structures for each model and attribution method.

To assess whether the attention was differentially allocated between structures, we focused on the contrast between arteries and veins. For each model and each attribution method, we computed paired-samples Cohen’s *d* to quantify the standardised difference between the RAI distributions for arteries and veins across the dataset^37^:

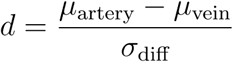

where *μ* denotes the mean RAI across all images, and *σ*_diff_ is the standard deviation of the RAI differences within each image. This paired effect size reflects how strongly attention differentiates artery-specific from vein-specific signal, and allows direct comparison across models.

### Control Model with Randomised Parameters

To assess whether the attribution maps produced by the different interpretability methods reflect learned model representations rather than trivial image-dependent artefacts, we used a control model with randomised parameters, a sanity check first proposed by Adebayo et al.^26^.

Specifically, we repeated the entire attribution-map extraction pipeline using a RETFound model with the same architecture and configuration as the fine-tuned models, but without loading any trained checkpoint. In this setting, all model parameters, including the transformer encoder and prediction head, remain at their random initialisation. Such random initialisation followed PyTorch’s default initialisation scheme for the corresponding layers, and to ensure reproducibility across folds, random seeds were fixed per fold prior to model instantiation. Apart from the absence of trained weights, all other aspects of the model configuration (architecture, input preprocessing, output dimensionality, and evaluation mode) were kept identical to those used for the fine-tuned models.

Attribution maps for the control model were generated using the same interpretability methods, visualisation targets, and post-processing steps as for the trained models. We then applied to these control-model attribution maps the same quantitative analyses as we did for the trained models, including vessel-structure overlap metrics and perturbation-based insertion and deletion tests.

## Notes

### Competing Interest Statement

The authors have declared no competing interest.

### Funding

- Swiss National Science Foundation: CRSII5_209510

